# A Genetic Method for Distinguishing Cryptic *Pocillopora* Species in French Polynesia without Sequencing

**DOI:** 10.64898/2026.04.25.720756

**Authors:** Francesca M. Cohn, Erika Johnston, Scott C. Burgess, Jordan A. Sims, Krithika Layagala, Pierrick Harnay, Hollie M. Putnam, Adrienne M.S. Correa

## Abstract

*Pocillopora* is a widespread, dominant reef-building coral genus in the Indo-Pacific that exhibits high morphological similarity and plasticity. Given this, genetic tools are needed to robustly identify *Pocillopora* individuals to the species level. Quick and accurate identification approaches for *Pocillopora* species are critical to estimating biodiversity patterns under current and future environmental challenges. In recent years, the mitochondrial open reading frame (mtORF) and a histone region (PocHistone) have been validated using genome-wide data to become the most widely used species-level markers for *Pocillopora*. However, Sanger sequencing of a large number of samples can be prohibitively expensive and sequencing facilities are not always readily available. Therefore, we present restriction fragment length polymorphism (RFLP) digests here that identify the six species of *Pocillopora* (*P. acuta*, *P.* cf. *effusa*, *P. grandis*, *P. meandrina*, *P. tuahiniensis*, and *P. verrucosa*) found in French Polynesia, without sequencing. In uninformed validation tests (*in silico* and *in vitro*), our protocol identified each *Pocillopora* species with 100% accuracy. Given their cost-effective, rapid nature, the tailoring of additional RFLP digest protocols to identify cryptic coral species in reef regions around the world will support foundational reef science, conservation and restoration initiatives.

## Introduction

*Pocillopora* (Lamarck 1816) is a genus of scleractinian reef-building corals widespread across reefs and reef zones in the greater Indo-Pacific. *Pocillopora* is notoriously plastic in its morphology (Todd 2008, Gélin et al., 2017; Johnston et al., 2018; Johnston et al., 2023; Marti-Puig et al., 2014; Paz-García et al., 2015; Pinzón et al., 2013, with environmental factors such as light intensity and water motion (Todd 2008; Mass and Genin, 2008; Edmunds and Burgess, 2025; Enriquez et al. 2017) contributing to overall skeletal morphology. However, despite gross morphological similarity (**Figure 1**), cryptic species differ in various ways, such as the corallivore grazing pressure they experience (Burgess et al. 2025), their reproductive timing (Harnay et al. 2025), Symbiodinaceae communities (Johnston et al., 2022a) and responses to temperature (Burgess et al. 2021) and light (Edmunds and Burgess 2025, Edmunds et al. *in review*). These differences contribute to a growing body of evidence that morphologically cryptic *Pocillopora* species do not necessarily share equivalent ecological niches in the reef system (Edmunds and Burgess, 2025; Johnston et al., 2022).

**Figure 1.**
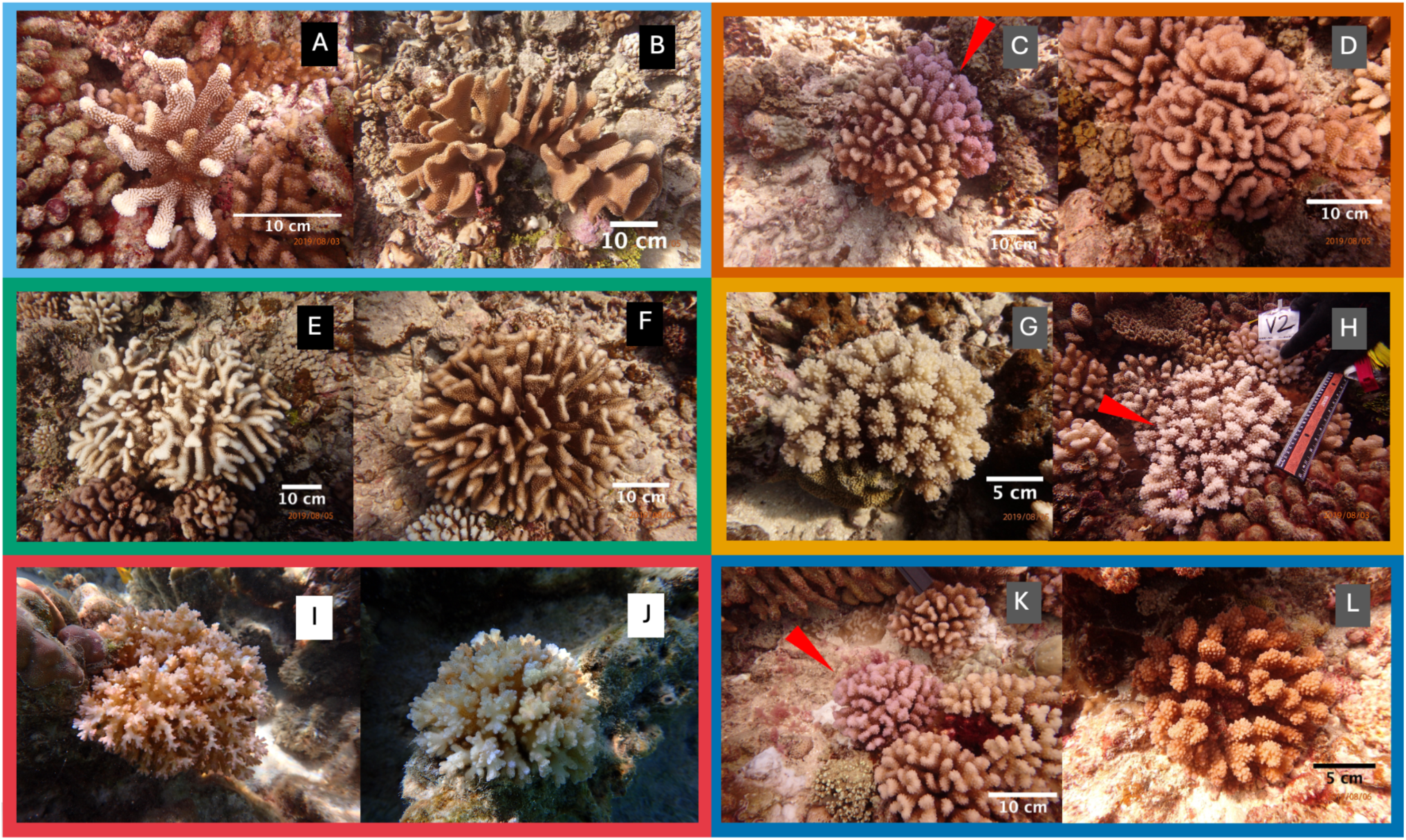
Pictures of cryptic *Pocillopora* species taken around Mo’orea, French Polynesia (A-L). Where multiple colonies are shown in a photo, the colony identified is shown with a red arrowhead. Species are as follows: A and B are *P. grandis*; C and D are *P. tuahiniensis*; E and F are *P.* cf. *effusa*; G and H are *P. verrucosa*; I and J are *P. acuta*; K and L are *P. meandrina*. Red arrows indicate the coral of interest where ambiguous. Individuals of the same species can look very different from each other (i.e., *P. grandis*, A and B), and individuals of different species can look very similar to each other (i.e., P*. tuahiniensis*, C and D; *P. verrucosa*, G and H; *P. meandrina*, K and L) due to phenotypic plasticity in this genus. The boxes behind letters A-L are color coded to group *Pocillopora* spp. that are easily confused with each other when they are adults: *P. grandis* and *P.* cf. *effusa* (black); *P. tuahiniensis*, *P. verrucosa*, and *P. meandrina* (grey). *P. acuta* (white) is the most reliably identifiable by morphology as an adult.

Furthermore, interspecific differences in bleaching support the value of *Pocillopora* spp. as a model for cryptic species resistance and resilience (natural and with human intervention) to climate change (Burgess, 2021; Burgess, 2025; Hughes et al., 2023). The ability to quickly and accurately identify cryptic *Pocillopora* species within a given morphotype (Oury et al., 2023, Voolstra et al., 2023, Oury et al. 2025) is therefore critical to estimating reef biodiversity and understanding how communities of morphologically indistinguishable species may contribute to the robustness of reefs against future environmental challenges (e.g., portfolio effects, Burgess et al. 2024, Edmunds and Burgess 2025).

To date, approaches using genetic markers (Flot et al., 2008; Pizón and Lajeunesse, 2011; Pizón, 2013; Schmidt-Roach et al., 2014; Marti-Puig et al., 2014; Gélin et al., 2017) have delineated species within *Pocillopora*, resolving phylogenetic relationships between historically confused types (**Table 1**; Johnston et al., 2022a; Oury et al., 2023). A standardized barcoding method using the mitochondrial open reading frame (mtORF, Flot et al., 2008; Pinzón, 2013; Johnston et al., 2018) and a histone region (PocHistone, Johnston et al. 2018) to delineate *Pocillopora meandrina* and *Pocillopora grandis*, which share mtORF haplotype 1, have in recent years become the most widely used species-level markers for *Pocillopora* (**Table 1**). Importantly, multiple independent genomic datasets and phylogenetic analyses have organized each mtORF haplotype and PocHistone variant into validated species, allowing researchers to identify species based on these markers, rather than whole genome sequencing (Johnston et al., 2022a; Oury et al., 2023; Voolstra et al., 2023). However, even Sanger sequencing the mtORF region may be prohibitively expensive for studies involving large sample sizes (Johnston et al., 2018). Sequencing is also frequently infeasible in remote locations where *Pocillopora* spp. occur. A restriction fragment length polymorphism (RFLP) digest and gel protocol was therefore developed to ameliorate these issues in Hawai’i (Johnston et al., 2018); this protocol supported reliable identification of *Pocillopora* spp. by leveraging interspecific nucleotide differences of the mtORF (Flot et al., 2008) and PocHistone regions (Johnston et al., 2018) without sequencing. Johnston et al. (2018) was the first to uniquely identify *Pocillopora meandrina* and *Pocillopora grandis*, which both share the same mtORF haplotype, haplotype 1, using a histone region.

**Table 1.** Table of naming conventions from prior studies and those used in this study for the species of *Pocillopora* found in and around Mo’orea, French Polynesia.

| Haplotype ID<br>Pinzon et al. (2013); Forsman et al. (2013); Burgess et al. (2021) | Lineage Current Name | Corresponding Lineages<br>Greek letters refer to Schmidt-Roach et al. (2014);<br>Roman numerals refer to Marti-Puig et al. (2014);<br>Genomic species hypotheses (GSH) refer to Oury et al. (2023) |
| --- | --- | --- |
| 2, 11 | *P. cf. effusa* | IIIa, GSH01 |
| 3a, 3b, 3c, 3d, 3e, 3f, 3h | *P. verrucosa* | γ, IIa, GSH13a, GSH13c, GSH13b |
| 10 | *P. tuahiniensis* | GSH14 |
| 5a | *P. acuta* | β, Ia, GSH05 |
| 1a, 1c, 1d, 1e, 8a, 9 | *P. meandrina* | m, IIb, GSH09b |
| 1a | *P. grandis* | e, IIb, GSH09c |

The restriction fragment length polymorphism (RFLP) protocol developed by Johnston et al. (2018) was designed to differentiate *Pocillopora* species in Hawai‘i (*Pocillopora acuta, Pocillopora damicornis, P. grandis, Pocillopora ligulata, P. meandrina*, and *Pocillopora verrucosa*). The complete protocol has not been validated with *Pocillopora* spp. in other geographic regions, which often harbor additional *Pocillopora* haplotypes and species (Oury et al. 2025, Connelly et al. 2026). Several enzymes that were formerly identified as sufficient to differentiate *Pocillopora* in Hawaiʻi are confounded by *de novo* mtORF haplotypes present at other locations due to the presence of different species. For example, the restriction enzyme *SacI-HF* does not cut haplotype 8a, a haplotype not present in Hawai‘i, but which is a variant of *P. meandrina* in French Polynesia (Johnston et al., 2018; Johnston et al., 2022a). Additionally, in *Pocillopora* samples from Hawai‘i (Johnston et al. 2018), the use of the restriction enzyme *AlwNI* differentiates *P. ligulata* (haplotype 6) from all other *Pocillopora* species, but this enzyme also cuts mtORF haplotype 11 (*Pocillopora* cf. *effusa*), which is not present in Hawai’i (Forsman et al., 2013; Gélin et al. 2017). However, in French Polynesia, *AlwNI* cannot be used to differentiate *P.* cf. *effusa* because *P.* cf. *effusa* consists of two mtORF haplotypes, haplotypes 2 and 11 (Johnston et al., 2022a), and *AlwNI* will not cut mtORF haplotype 2. RFLP protocols that accurately discern cryptic *Pocillopora* species in other regions of the Pacific (besides Hawai’i, Johnston et al., 2018) have not been available to date because extensive knowledge of existing *Pocillopora* species in the region of interest a prerequisite for RFLP protocol development.

Here, we describe an RFLP protocol that uses previously established mtORF and PocHistone regions to identify species of *Pocillopora* in French Polynesia, broadening the geographic applicability of RFLP methods for cryptic species identification. The diversity of *Pocillopora* species at Mo’orea has been extensively documented (Forsman et al., 2013; Johnston et al., 2022a; Johnston et al., 2022b; Burgess et al., 2024), and therefore RFLP methods could be developed for regionally specific mtORF haplotypes and species (e.g. *P.* cf. *effusa*: haplotypes 2, 11; *P. meandrina* haplotype 8a; *P. tuahiniensis*: haplotype 10). Additionally, *Pocillopora* are a dominant reef building genus in this region (Burgess et al., 2021; Burgess et al., 2024; Edmunds and Burgess, 2025; Burgess et al., 2025; Johnston et al., 2022b) contributing to ecological recovery (Tsounis and Edmunds, 2016; Burgess et al., 2021) following disturbance events including bleaching events, cyclones, and crown of thorns sea star outbreaks (Pratchett et al., 2011; Pratchett et al., 2013; Edmunds et al., 2016). Much of this research, however, has largely documented trends at the genus level (e.g., Pratchett et al., 2013, Speare et al., 2024) because species identification of *Pocillopora* using morphology is unreliable. We anticipate that the relatively inexpensive, rapid and sequencing-independent protocol presented here for the identification of cryptic *Pocillopora* species will facilitate species level monitoring and research efforts on French Polynesian reefs.

## Methods

Mo’orean reefs harbor broadly distributed, common *Pocillopora* species, as well as several endemic mtORF haplotypes (following the delineation and nomenclature established by Pinzón et al. 2013; Forsman et al. 2013). There are six species, containing 17 associated mtORF haplotypes, that are common on reefs around Mo’orea (Edmunds, 2016, 2014; Burgess et al., 2021, 2024, 2025; Johnston et al., 2022a; Johnston and Burgess, 2023): *P. acuta*: 5a (Rouzé et al., 2019; Burgess et al., 2024); *P.* cf. *effusa*: 2, 11 (Burgess et al., 2021); *P. grandis*: 1a (Johnston et al., 2022a; Johnston et al., 2022b; Burgess et al., 2025); *P. meandrina*: 1a (same mtORF but different PocHis amplicon as *P. grandis*), 1c, 1d, 1e, 8a, 9 (Burgess et al., 2021; Supp. Material); *P. tuahiniensis:* 10 (Johnston et al., 2022a; Burgess et al., 2025; Johnston and Burgess et al., 2023); and *P. verrucosa*: 3a, 3b, 3c, 3d, 3e, 3f, 3h (**Table 1**; Burgess et al., 2025). *Pocillopora damicornis* and *Pocillopora woodjonesi* have also been reported at low abundance based on gross morphology in the Society Islands (Edmunds et al., 2016), and *Pocillopora damicornis* was reported at low levels in the Southern and Cook Islands based on mitochondrial ATP synthase subunit 6 (atp6) and putative control region (pmapcr, Mayfield et al., 2015). However, given the lack of genetic detection of *P. damicornis* and *P. woodjonesi* using widely applied genetic approaches (mtORF and PocHistone) despite extensive studies in Mo’orea (Burgess et al., 2021, 2024, 2025; Johnston et al., 2022b; Johnston and Burgess 2023, Harnay et al. 2025), these putative species were not included in the RFLP protocol presented here.

To develop a RFLP-based species identification protocol for *Pocillopora* species in Mo’orea and the greater French Polynesian region, we used an iterative process of: (1) identifying species-defining single nucleotide polymorphisms (SNPs) in the mtORF and PocHistone regions (Johnston et al. 2022) and *in silico* identification of restriction enzymes that target these sites; and (2) testing the utility of candidate restriction enzymes via *in vitro* digestion of DNA from previously sequenced *Pocillopora* samples. The digestion protocol was optimized based on the ability of its component restriction enzymes to explicitly distinguish *Pocillopora* spp. and haplotypes based on amplicon band sizes. Lastly, *in silico* and *in vitro* uninformed validation tests (i.e., species IDs were temporarily redacted from samples) were run on sample sets to ensure accuracy and usability of the finalized enzyme protocol.

### In Silico SNP and Enzyme Identification Methods

Species-specific *Pocillopora* single nucleotide polymorphisms (SNPs) were identified from a *Geneious Prime* (v.2024.0.02) alignment of the 17 unique mtORF haplotypes (1a, 1c, 1d, 1e, 2, 3a, 3b, 3c, 3d, 3e, 3f, 3h, 5a, 8a, 9, 10, 11; **Supplemental Data S1**) documented from Mo’orea based on ∼4,500 samples drawn from Johnston et al. (2022b), Burgess et al. (2021), Burgess et al. (2024), Burgess et al. (unpublished), and Harnay et al. (unpublished). Candidate restriction enzymes were tested *in silico* using the *Restriction Analysis* feature of *Geneious Prime*. Restriction enzymes were selected for testing and validation based on their potential to differentiate two or more of the focal *Pocillopora* species. A similar selection process was attempted using an alignment of the PocHistone amplicon for haplotypes associated with each focal *Pocillopora* species based on a total of 41 samples (Johnston et al., 2018, **Supplemental Data S2**). However, high intra-haplotype variability in the PocHistone amplicon limited this approach as exemplified by the attempted *HgaI* digest of PocHistone described below (**Supplemental Figure S1**). Therefore, the PocHistone amplicons were only used as previously described in Johnston et al. (2018) in a *XhoI* digest to differentiate *P. grandis* and *P. meandrina*. To determine the location of each enzyme’s cut sites and all expected band sizes of the cut mtORF amplicon, the Flot et al. (2008) primers were mapped onto the *P. grandis* mitochondrial genome (GenBank accession NC_009798.1). This reference genome was added to the mtORF alignment to determine where predicted cut sites fell within the mtORF amplicon.

### In Vitro Enzyme Testing Methods

A total of eight candidate enzymes were identified for further testing *in silico*: *AciI* (New England Biolabs (NEB), MA, USA, #R0551S), *BanI* (NEB, #R0118S), *BceAI* (NEB, #R0623S), *BseYI* (NEB, #R0635S), *EcoRV-HF* (NEB, #R3195S), *HgaI* (NEB, #R0154S), *NlaIV* (NEB, #R0126S), *SacI-HF* (NEB, #R3156S), and *XhoI* (NEB, #R0146S). Each enzyme was used in *in vitro* restriction digests applied to mtORF amplicons generated from 42 samples of *Pocillopora* spp. representing 12 haplotypes (1a, 2, 3a, 3b, 3e, 3f, 3h, 5a, 8a, 10, 9, 11), collected from 5, 10, and 20m depth at four sites across the forereef, lagoon, and fringing reef zones of the island of Mo’orea (Johnston et al., 2022b; Burgess et al., 2021, 2024; Burgess et al., unpublished). Additionally, the associated PocHistone amplicon of each of the above samples was tested with *XhoI* to test the digest’s ability to differentiate *P. grandis* and *P. meandrina* species among haplotypes from French Polynesia. The utility of restriction enzyme digests in accurately parsing *Pocillopora* species was assessed based on: (1) the extent to which the RFLP ‘fingerprints’ produced by an enzyme matched the band size expected based on *in silico* analysis of sequence data; and (2) the ability of human users to accurately visually distinguish distinct banding patterns in gels.

In preparation for restriction enzyme digest tests, 50µl aliquots of mtORF amplicons were generated from DNA extractions of the 42 *Pocillopora* samples using primers FatP6.1 (5′-TTTGGGSATTCGTTTAGCAG-3′) and RORF (5′-SCCAATATGTTAAACASCATGTCA-3′) (Flot et al. 2008). Polymerase chain reactions (PCRs) were run using Taq PCR Master Mix Kits (Qiagen, Hilden, Germany) and contained 25µl of master mix, 22µl of H_2_O, 1µl of each primer (forward and reverse, 10uM concentration), and 1µl of sample DNA (at approximately a 20-30 ng/µl concentration). The thermocycler protocol included initial denaturation at 94°C for 60 sec; 40 cycles of 94°C for 30 sec, 53°C for 30 sec, and 72°C for 75 sec; and a final extension of 72°C for 5 min (Flot et al. 2008, Johnston et al. 2018). PocHistone amplicons were also generated from DNA from each *Pocillopora* sample using PocHistoneF (5′-SCCAATATGTTAAACASCATGTCA-3′) and PocHistoneR primers (5′-TATCTTCGAACAGACCCACCAAAT-3, Johnston et al. 2018) using the same reagents and reaction volumes detailed above for mtORF reactions. However, the following thermocycler program was used in PocHistone PCR reactions: initial denaturation at 94°C for 60 sec; 40 cycles of 94°C for 30 sec, 53°C for 30 sec, and 72°C for 1 min; and a final extension of 72°C for 5 min. Amplicons were stored at −20°C prior to undergoing restriction enzyme digestion.

### Restriction Enzyme Digestion Conditions for In Vitro Testing

Generally, to perform enzyme digestions, 5µL of PCR amplicons generated from each *Pocillopora* sp. were digested with 0.5µL of digestion buffer and 0.5µL of one restriction enzyme or 1µL of two combined restriction enzymes for 1 hour at 37°C, followed by a 20 minute enzyme denaturation step. Reaction reagents and denaturation temperatures varied for each restriction enzyme assay and are detailed in **Table 2**. Following digestion, 5µL of digested PCR amplicons were run on a 2% agarose gel at 100V for 45 minutes. Length of DNA fragments produced was determined by comparison to a 50bp DNA ladder (NEB, #N3236). The restriction enzyme *NlaIV* was previously validated for grouping *P. damicornis* and *P. acuta* in an RFLP protocol for Hawai‘i (Johnston et al. 2018) and was not included in the *in vitro* enzyme identification process; this enzyme was included, however, in the *in vitro* uninformed validation.

**Table 2.**
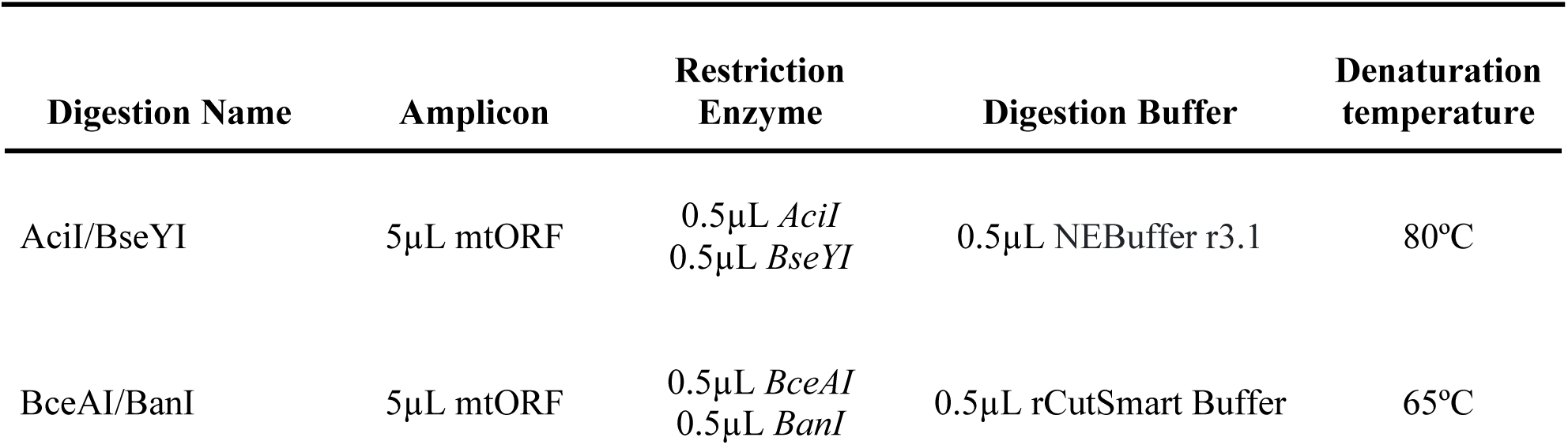
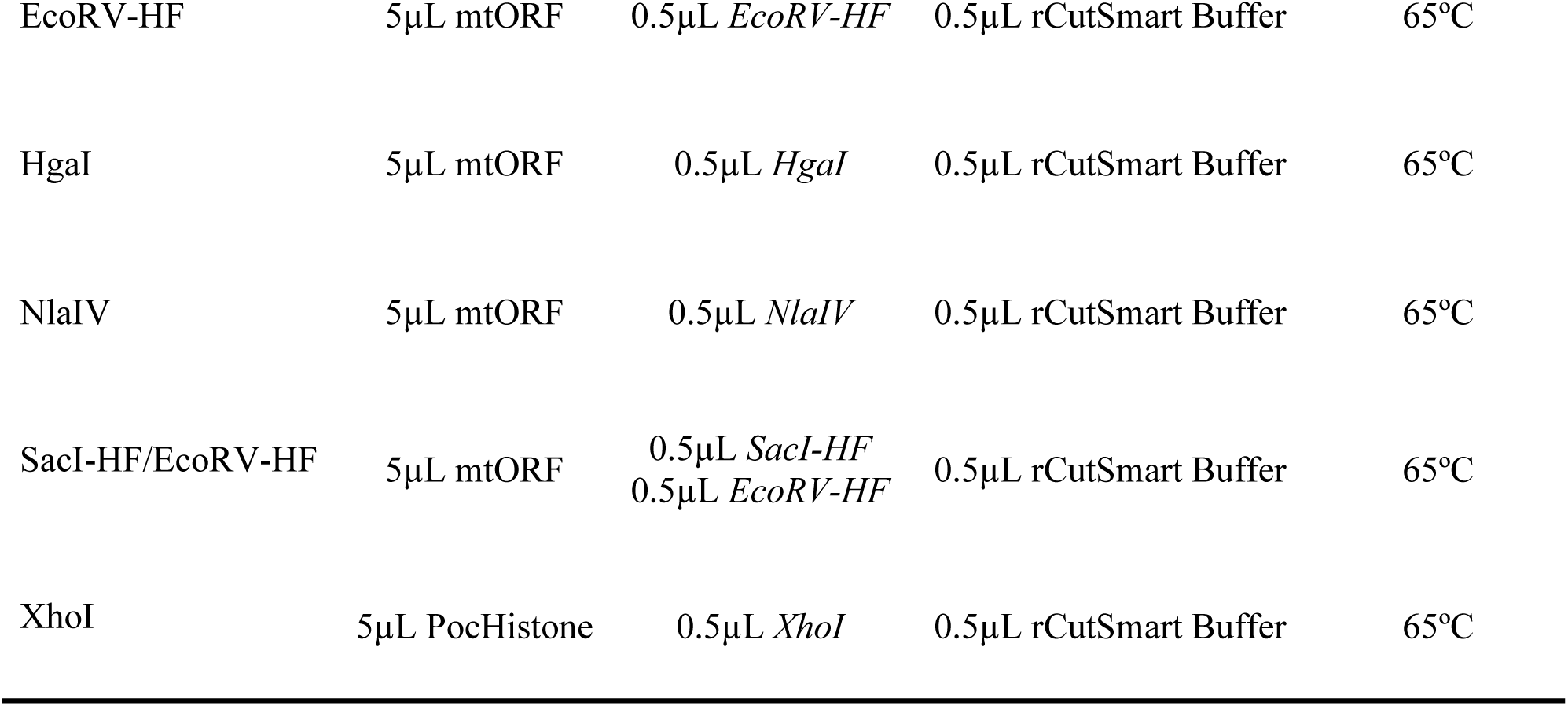
Reagent volumes and denaturation temperatures for all restriction enzyme digestions tested.

### In Silico and In Vitro Uninformed Validation Methods

*In silico* uninformed validations of *BseYI/AciI*, *EcoRV-HF*, and *NlaIV* were next performed on an mtORF haplotype dataset generated from 673 *Pocillopora* samples collected in French Polynesia (Johnston et al., 2022b), a subset of the ∼4,500 samples used to construct the alignment used for the *In Silico SNP and Enzyme Identification Methods*. Using the *Restriction Analysis* feature of Geneious Prime 2024.0.02, mtORF haplotypes were digested with *BseYI*/*AciI,* and each haplotype was categorized as either *P. verrucosa*, *P. tuahiniensis*, or one of the other four *Pocillopora* species. A *Restriction Analysis* digest was then emulated with *EcoRV-HF*, and each haplotype was categorized as either *P. cf. effusa* or one of the other five *Pocillopora* species. A final *Restriction Analysis* digest was then run with *NlaIV*, and each haplotype was categorized as (1) *P. tuahiniensis* or *P. verrucosa*, (2) *P. grandis*, *P. meandrina*, or *P. cf. effusa*, or (3) *P. acuta.* The predicted species for each sample was then compared against the sample metadata to confirm that the correct species had been predicted by the *in silico* analyses. *XhoI* was not included in the uninformed test because PocHistone sequences were not available from Johnston et al. (2021). However, *XhoI* cutting of PocHistone amplicons was extensively shown to distinguish *P. grandis* and *P. meandrina* by Johnston et al. (2018) and it was established that *XhoI* digested the *P. grandis* haplotype, while leaving all *P. meandrina* haplotypes uncut (including the endemic 8a haplotype) during *in silico* enzyme testing (Results).

An *in vitro* uninformed validation was then conducted on 36 samples of *Pocillopora*, representing eight haplotypes from *P. acuta* (haplotype 5a, n=5 from Mo’orea and n=1 from URI aquarium), *P. meandrina* (haplotypes 1a, 8a, n=5 each), *P. cf. effusa* (haplotype 11, n=5), *P. grandis* (haplotype 1a, n=5), *P. tuahiniensis* (haplotype 10, n=5), and *P. verrucosa* (haplotypes 3a, n=1; 3b, n=4) sampled at 2m depth from the lagoonal reefs of the north shore of Mo’orea, French Polynesia (**Supplemental Figure S2, Supplemental Table S1**). Initial species identifications were performed at the University of Rhode Island by Sanger sequencing the mtORF region using FatP6.1 and RORF primers (**Supplemental Data S3**). Samples with haplotype 1 were then amplified using PocHistone primers, digested with *XhoI*, and visualized as described above for species-level identification. DNA aliquots of each sample (with species IDs redacted) were sent to the University of California Berkeley, where the uninformed validation of this protocol was performed. PocHistone and mtORF amplicons were generated and each of four digests with an additional alternative *BceAI/BanI* digest (**Supplemental Materials and Methods: *BanI/BceAI Digest*)** was run according to the methods described in ***In Vitro Enzyme Identification***.

## Results

### In Silico SNP and Enzyme Identification

*P. acuta* (haplotype 5a) was differentiated from all other *Pocillopora* species in Mo’orea via mtORF using a previously identified 341 bp fixed Guanine SNP recognized by *NlaIV* and validated in Johnston et al. (2018). This polymorphism additionally distinguished *P. meandrina*, *P. grandis*, and *P. cf. effusa* from all other species for further differentiation by a fixed guanine SNP at 312 bp in *P. cf. effusa*, *P. acuta*, *P. meandrina*, and *P. grandis*. All species share a cut site for this restriction enzyme at 514 and 515 bp (**Table 3**).

**Table 3.** Summary of *Pocillopora* species found in Mo’orea, French Polynesia, their associated haplotypes, the amplicon and restriction enzyme (RE) used for identification, resulting band fragment size of that species, and resulting band fragment size of other species when cut with that restriction enzyme. Additional digests required to fully identify each *Pocillopora* species are listed in subsequent columns.

| Species | Haplotypes | Amplicon | RE | Fragment size (bp) | Cuts other Species | Additional Digests | Amplicon | RE | Fragment size (bp) | Cuts other Species |
| --- | --- | --- | --- | --- | --- | --- | --- | --- | --- | --- |
| *P. verrucosa* | 3a, 3b, 3c, 3d, 3e, 3f, 3h | mtORF | (1) BseYI<br>C'CCAGC<br>GGGTC'G<br>AciI<br>C'CGC<br>GGC'G | 209, 338, 430 | All other species<br>430, 546 | No |  |  |  |  |
| *P. tuahiniensis* | 10 | mtORF | (1) BseYI<br>C'CCAGC<br>GGGTC'G<br>AciI<br>C'CGC<br>GGC'G | 145, 186, 209,<br>430 | All other species<br>430, 546 | No |  |  |  |  |
| *P. cf. effusa* | 2, 11 | mtORF | (1) EcoRV -<br>HF<br>GAT'ATC<br>CTA'TAG | 339, 639 | All other species<br>978 | No |  |  |  |  |
| *P. acuta* | 5a | mtORF | (1) NlaIV<br>GGN'NCC<br>CCN'NGG | 30, 171, 313, 463 | mean, grandis, cf.<br>effusa<br>202, 313, 463<br>verr, tuah<br>463, 515 | No |  |  |  |  |
| *P. meandrina* | 1a, 1c, 1d, 1e, 8a, 9 | mtORF | (1) NlaIV<br>GGN'NCC<br>CCN'NGG | 202, 313, 463 | acuta<br>30, 172, 313, 463<br>verr, tuah<br>462, 515<br>cf. effusa<br>202, 313, 463 | (2) EcoRV-HF<br>to remove P. cf.<br>effusa | (2)<br>PocHistone | (3) XhoI<br>C'TCGAG<br>GAGCT'C | 287, 382<br>(Johnston et al<br>2018) | All other species<br>669 |
| *P. grandis* | 1a | mtORF | (1) NlaIV<br>GGN'NCC<br>CCN'NGG | 202, 313, 463 | acuta<br>30, 172, 313, 463<br>verr, tuah<br>462, 515<br>cf. effusa<br>202, 313, 463 | (2) EcoRV-HF<br>to remove P. cf.<br>effusa | (2)<br>PocHistone | (3) XhoI<br>C'TCGAG<br>GAGCT'C | 669 (Johnston et<br>al 2018) | All other species<br>669 |

To differentiate *P. cf. effusa* (haplotype 2, 11) from all other species, a fixed cytosine SNP at 342 bp was identified as a cut site of the restriction enzyme *EcoRV-HF* in the multispecies mtORF alignment. This fixed cytosine SNP at 342 bp in *P. cf. effusa* is not shared with any other species (**Table 3**).

*P. verrucosa* (haplotypes 3a, 3b, 3c, 3d, 3e, 3f, 3h) and *P. tuahiniensis* (haplotype 10) were differentiated from each other and all other species using a co-digest of *AciI* and *BseYI* targeting SNPs in the mtORF alignment. To differentiate *P. verrucosa* from all other species, a 210 bp fixed cytosine SNP was identified in *P. verrucosa* and *P. tuahiniensis* and is recognized by *AciI.* The same restriction cut site is shared by all species at 546 bp. To simultaneously differentiate *P. tuahiniensis* from all other species, a fixed cytosine SNP is used at 398 bp and recognized by *BseyI* (**Table 3**).

*P. grandis* and *P. meandrina* are easily confused at small colony sizes and share the 1a mtORF haplotype. The Johnston et al. (2018) protocol uses a *XhoI* digest of the PocHistone amplicon to differentiate these species, as *XhoI* cuts *P. grandis*, but not *P. meandrina,* due to a fixed thymine polymorphism at 279 bp (**Table 3**). In the process of designing this protocol, it was confirmed that the *XhoI* digest of *P. grandis* amplicons may produce either 2 or 3 bands when run out on a gel. Further sequencing (restriction site associated DNA sequencing, RADseq) of *Pocillopora* samples from Mo’orea has revealed that some *P. grandis* individuals are heterozygotic for the *XhoI* restriction cut site (5’C−TCGAG3’), suggesting that some individuals may be monozygotic for absence of the cut site. Of 194 samples collected from Mo’orea that were identified as *P. meandrina* by sequencing the mtORF region followed by the PocHistone RFLP protocol (i.e., these 194 samples were monozygotic for absence of the *XhoI* cut site), only one sample out of 194 (0.5% of colonies analyzed), was erroneously identified as *P. grandis* following the analysis of RADseq data (**Supplemental Materials and Methods: *RADseq Histone Analysis***).

### In Vitro Enzyme Testing

*EcoRV-HF* (mtORF), *BseYI/AciI* (mtORF), *NlaIV* (mtORF), and *XhoI* (PocHistone) were selected for this protocol because they produced RFLP ‘fingerprints’ that matched the band sizes expected based on *in silico* analysis of sequence data and were easily distinguishable visually on gels (**Figure 2**). Other enzymes, such as *SacI-HF* and *HgaI* were removed because all the haplotypes of a *Pocillopora* species did not have the same banding pattern (**Supplemental Figure S1, S3**). While other enzymes were possible candidates for the final protocol, including *BceAI/BanI* to identify *P.* cf. *effusa* and group together *P. grandis* (Haplotype 1), *P. meandrina* (Haplotype 1a, 1c, 1d, 1e, 8a, 9), and *P. acuta* (Haplotype 5a), these were not selected for the protocol because they increased the number of enzymes required to identify all *Pocillopora* species sequentially (**Supplemental Materials and Methods: *BanI/BceAI Digest*, Supplemental Figure S4, Supplemental Table S2)**. The RFLP digests described above are combinatorially able to rapidly and unambiguously identify the six *Pocillopora* species present in Mo’orea. For the majority of *Pocillopora* samples, only one digest was needed to identify the corresponding species after the sample had been amplified at the mtORF or PocHistone regions. The *EcoRV-HF* digest identified *P. cf. effusa* (FatP6.1 and RORF primers); the *NlaIV* digest identified *P. acuta* (FatP6.1 and RORF primers); the *BseYI/AciI* digest identified both *P. tuahiniensis* or *P. verrucosa* (FatP6.1 and RORF primers); the *XhoI* digest identified *P. grandis* (PocHistoneF and PocHistoneR primers). Three digests and both primer sets are required to identify *P. meandrina*: (*NlaIV* (mtORF), *EcoRV-HF* (mtORF), and *XhoI* (PocHistone), **Table 3**). Using these enzymes, an unknown *Pocillopora* sample from French Polynesia can be identified to species using a sequential protocol of up to six steps (***Finalized Sequential Protocol***).

**Figure 2.**
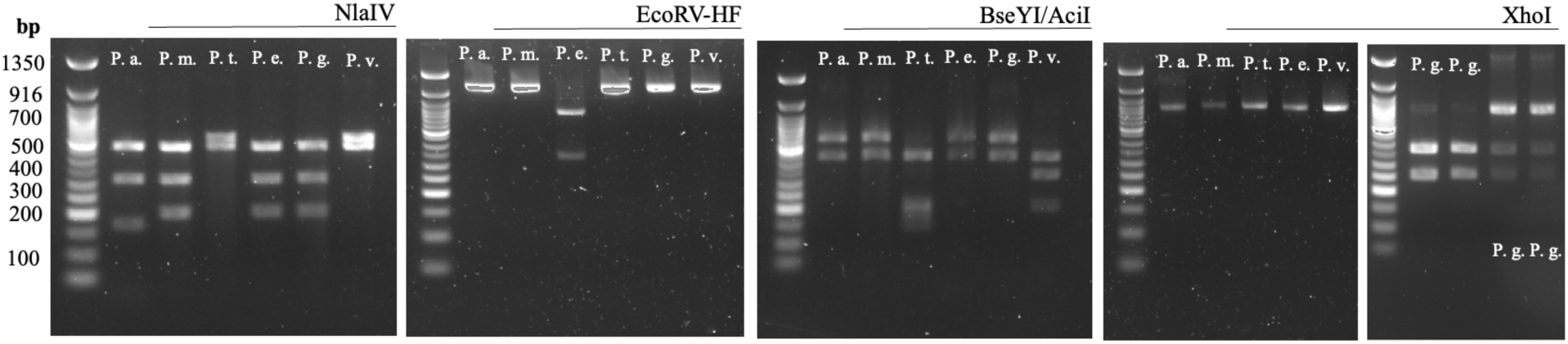
Gel band patterns generated by each restriction enzyme digest, which identify species of *Pocillopora* found in French Polynesia: *Pocillopora acuta* (P.a.), *Pocillopora meandrina* (P.m.), *Pocillopora tuahiniensis* (P.t.), *Pocillopora* cf. *effusa* (P.e.)., *Pocillopora grandis* (P.g.), *Pocillopora verrucosa* (P.v.), Left to right: *NlaIV* digest of the MtORF amplicon for all six *Pocillopora* species; *EcoRV-HF* digest of the MtORF amplicon for all six *Pocillopora* species; *BseYI/AciI* digest of the MtORF amplicon for all six *Pocillopora* species; *XhoI* digest of the PocHistone amplicon for all six *Pocillopora* species. The *P. grandis* digests are two different samples, representative of the (left to right) 2 band vs. 3 band cut patterns.

### In Silico and In Vitro Uninformed Validation of the Protocol

The *in silico* uninformed validation of *BseYI/AciI*, *EcoRV-HF*, and *NlaIV* digestions identified 100% of the samples (n = 673) accurately based on each digest. Additionally, the *in vitro* uninformed validation identified 100% of samples (n=36) to species accurately by RFLP, as confirmed by prior (uninformed) sequencing the mtORF region of each sample (**Figure 3**, **Supplemental Data S3**).

**Figure 3.**
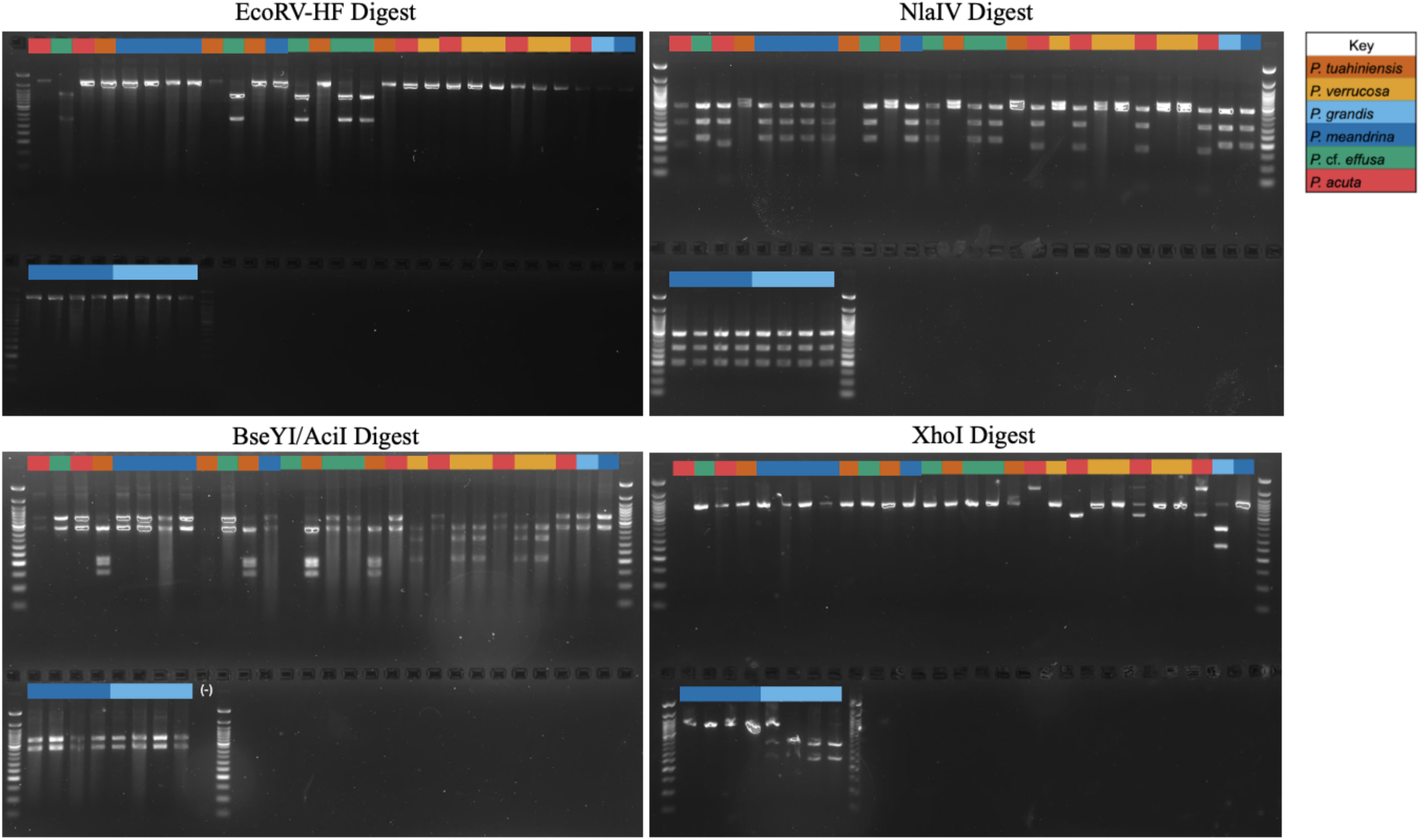
Results of the *in vitro* uninformed validation for each digest run on 36 unidentified *Pocillopora* samples representing *P. acuta*, *P. meandrina*, *P.* cf*. effusa*, *P. grandis*, *P. tuahiniensis*, and *P. verrucosa* collected in French Polynesia. Lanes 1, 9, and 13 did not appear in some digests but were still identifiable from running all 4 digests. 100% of samples were identified accurately.

### Finalized Sequential Protocol

For a collection of *Pocillopora* samples with no prior information about their species identity, the sequential protocol is (**Figure 4**):

1. PCR amplify the mtORF region of each sample using the FatP6.1 and RORF primers.
2. Digest the mtORF amplicons with the restriction enzyme *NlaIV* to (1) distinguish *P. acuta* from all other species, (2) group *P. tuahiniensis* and *P. verrucosa*, and (3) group *P. effusa* + *P. meandrina* + *P. grandis*)
3. For samples that are in the *P. tuahiniensis*/*P. verrucosa* group, digest the mtORF amplicon with the restriction enzymes *BseYI and AciI* to distinguish *P. tuahiniensis* and *P. verrucosa*.
4. For samples that are in the *P. effusa*/*P. meandrina*/*P. grandis* group, digest the mtORF amplicon with the restriction enzyme *EcoRV-HF* to distinguish *P. effusa* from *P. meandrina* and *P. grandis*.
5. For samples that are in the *P. meandrina* + *P. grandis* group, PCR amplify the PocHistone amplicon using the PocHistoneF and PocHistoneR primers.
6. Digest the PocHistone amplicons with *XhoI* to distinguish *P. meandrina* and *P. grandis*.

**Figure 4.**
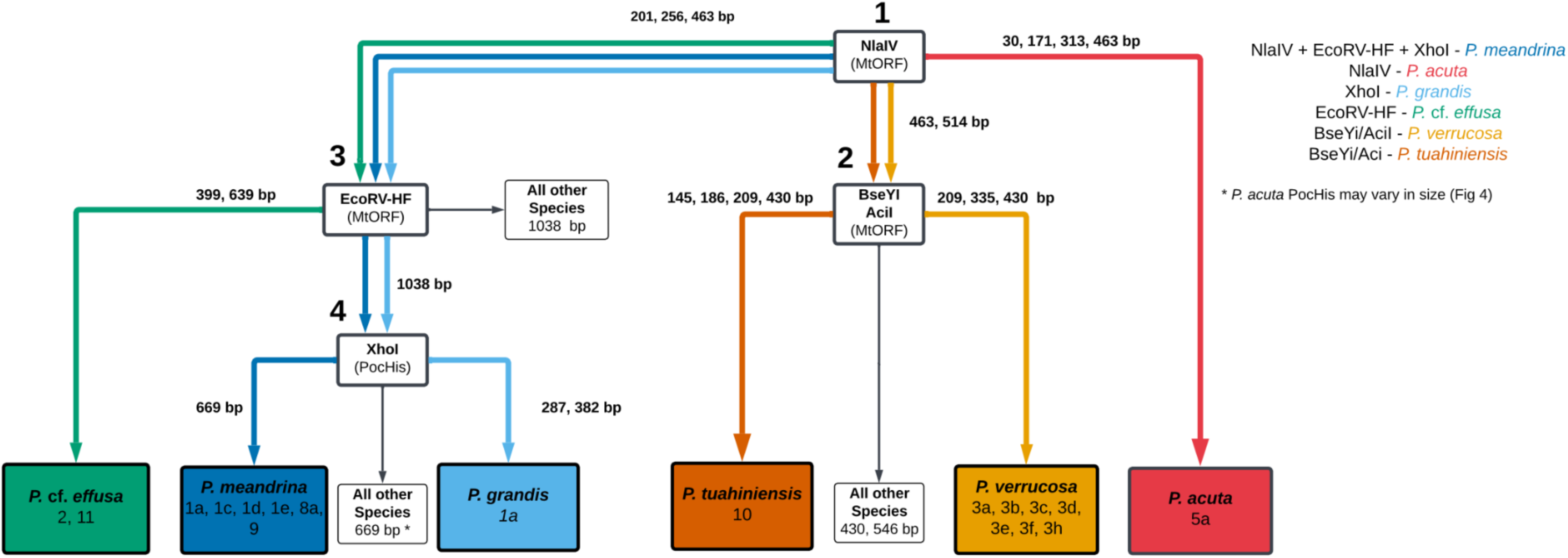
Flow chart of digests required for the identification of each species of *Pocillopora* present in French Polynesia. Colored arrows correspond to the path taken through the diagram for each species. The large numbers indicate the order the digests should be performed in as part of the Finalized Sequential Protocol to identify all species with no prior information.

## Discussion

This study presents a restriction enzyme digest method to rapidly and unambiguously differentiate the six species of cryptic *Pocillopora* present in Mo’orea, French Polynesia, based on their mtORF and PocHistone haplotypes. Taken together with the methods described in Johnston et al. (2018), this study indicates that the development of additional region-specific, rapid, and sequencing-free RFLP protocols holds merit for cryptic coral species identification in geographic reef regions (e.g., the wider Pacific and Red Sea for *Pocillopora*).

In Mo’orea, the morphological similarity and plasticity of *Pocillopora* species have historically posed a major roadblock to species-specific research (**Figure 1**; Johnston et al., 2022a, 2023; Burgess et al., 2024, 2025). For example, *P. tuahiniensis, P. verrucosa,* and *P. meandrina* share similar gross skeletal morphologies (Johnston et al., 2022b, 2023; Burgess et al., 2024) and an overlapping depth distribution (**Figure 1**; 2–20m; Burgess et al., 2024). The method presented here provides a single digest of the mtORF amplicon with *BseYi/AciI* to identify *P. verrucosa* and *P. tuahiniensis*, and distinguish between them (**Table 3**)*. P. meandrina* and *P. grandis* are also often difficult to distinguish at smaller sizes, and both can also be confused with *P. verrucosa* (Burgess et al. 2025)*. XhoI* digest of the PocHistone amplicon identifies *P. grandis* samples; additional digests of mtORF amplicons with *NlaIV*, *EcoRV-HF*, and *BseYi/AciI*, respectively are required to distinguish *P. meandrina* from *P. verrucosa* (**Table 3**). Finally, adult *P. grandis* and *P.* cf. *effusa* can attain large colony sizes and can be morphologically indistinguishable (**Figure 1**; Johnston et al., 2022b). These species can be separated with a single digest of the mtORF amplicon with *EcoRV-HF* to identify *P.* cf. *effusa* samples, followed by an additional *XhoI* digest of the PocHistone amplicon to identify *P. grandis* (**Table 3**). As suggested by these examples, the methods presented in this paper can be used individually to identify specific target species, or combined to identify all species, making the protocol highly customizable to the identification needs of any project involving French Polynesian *Pocillopora* species. However, in most cases (especially for coral recruits), the finalized sequential protocol (**Figure 4**) must be run to accurately identify all *Pocillopora* species by process of elimination.

The biggest advantage of the method presented here is its application to remote field sites, where CITES (Convention on International Trade in Endangered Species of Wild Fauna and Flora) permitting can make the export of coral samples difficult. Many of the islands and atolls surrounding Mo’orea are ecologically salient locations for coral research because their remoteness and small human populations present relatively less disturbed environments for documenting baseline reef dynamics (Wright et al., 2025). The remoteness that contributes to the desirability of these locations also presents challenges for molecular work and highlights the need for field-adaptable genetic identification protocols such as the one we present here. With the advent of small, modular, and open-source PCR technology (e.g., prebuilt kits or DIY arduino or raspberry pi-based thermocyclers), the methods described in this paper allow for fast and efficient confirmation of *Pocillopora* species at remote field sites without access to sequencing facilities (see example protocols at Cohn et al., 2026, Edmunds et al., *in review*).

This RFLP protocol provides a cost effective alternative to the amplicon Sanger sequencing that was previously required to resolve morphologically similar *Pocillopora* spp. (Johnston et al. 2018). At the time of publication, the enzymes and associated buffers used in these methods cost approximately $4.20 USD per sample to run all four digests without any enzyme dilutions – roughly one-third of Sanger sequencing costs ($7.00 USD total to sequence in both directions). Additionally, depending upon the concentration of amplicons, 1:1 enzyme dilutions in water may be used to reduce RFLP protocol costs by half (to $2.10 USD per sample). Therefore, the RFLP protocol provides a flexible range of pricing that depends on the enzymes required, as opposed to a fixed sequencing rate that may be unnecessarily costly (i.e., when trying to distinguish between only 2 species, such as *P. verrucosa* and *P. tuahiniensis*).

We detected variation in the PocHistone amplicon used to differentiate *P. grandis* from *P. meandrina*. Unlike the mtORF amplicon, a mitochondrial marker, the PocHistone amplicons are generated from nuclear DNA and may therefore be heterozygous. We found that *P. grandis* amplicons in French Polynesia may produce either two or three bands following digestion, suggesting that some of the population is heterozygous for the *XhoI* cut site (5’ C/TCGAG 3’; **Figure 2**). While this does not interfere with identification of *P. grandis* using our RFLP protocol, this finding indicates that there may be *P. grandis* colonies that are homozygous for the absence of the *XhoI* cut site. However, evidence that variation in this cut site leading to the enzyme not recognizing the site (i.e., C/TCGCG as is observed in *P. meandrina*) in *P. grandis* was very rare – of the 194 colonies identified as either *P. meandrina* or *P. grandis*, only one (0.5%) of the *P. grandis* colonies was homozygous for the *XhoI* cut site absence (C/TCGCG) (**Supplemental Materials and Methods: *RADseq Histone Analysis***).

We also found a PocHistone amplicon anomaly among *P. acuta* samples. This included a 345 bp indel starting at position 538 bp (identified by Sanger sequencing; GenBank accession XXXX), resulting in a substantially longer PocHistone amplicon (∼1000 bp) compared to the 669 bp observed in other species (**Supplemental Figure S5**). Additionally, another *P. acuta* anomaly exists where another band ∼500bp is present, sometimes with the 1014 bp indel band and expected 669 bp band. However, this does not impact the ability to diagnose *P. grandis* with the *XhoI* digest, as only *P. grandis* presents the diagnostic 287 bp and 382 bp bands when digested (**Supplemental Figure S6**). Despite these caveats, the uninformed validation used in this study was able to identify *Pocillopora* samples (n=36) to species with 100% accuracy (**Figure 3**).

In terms of mtORF variation, it is noteworthy that in French Polynesia, *P. verrucosa* is represented by a large number of haplotypes (3a, 3b, 3c, 3d, 3e, 3f, 3h), and many of these haplotypes have few SNP differences between them (e.g. 3b and 3d; 3h and 3e: one SNP difference; 3a and 3b: two SNP differences). In Mo’orea, these haplotypes all correspond to *P. verrucosa* (Johnston et al., 2022a). However, in some study locations outside of French Polynesia, some of these haplotypes represent distinct species. For example, mtORF haplotype 3a and 3d are called *Pocillopora lacera* in the Eastern Tropical Pacific (Connelly et al., 2025; Oury et al., 2023; Oury et al., 2024) and haplotype 3 is called *Pocillopora favosa* in the Arabian Peninsula (Oury et al., 2025).

For *Pocillopora* spp., beyond Mo’orea, mtORF haplotypes have been documented at neighboring islands, including Tetiaroa, Tahiti, and Maiao (Edmunds et al., 2016), with no new haplotypes identified at these locations. The Tara Pacific expedition (2016–2018) sampled colonies resembling a typical *P. meandrina* morphology spanning 18,000 km of the Pacific Ocean, including Mo’orea, Tenoko, Tekava, and Kamaka from the Gambiers, and Aitutaki from the Cook Islands. Samples of *Pocillopora* spp. from the expedition did not identify any haplotypes unaccounted for in our methods, further supporting the applicability of our method to the larger region of French Polynesia. However, we do caveat that study sites located further from Mo’orea have the potential to contain species or haplotypes not represented in our results, which may reduce the accuracy of the RFLP protocol presented here at those locations. For example, *P. damicornis*, identified using variable sites with the mitochondrial ATP synthase subunit 6 (*atp6*) and putative control region (*pmapcr*), has been found in low abundance in the Southern and Cook Islands of French Polynesia and is not included in these methods (Mayfield et al., 2015).

However, if applying the RFLP protocol here to the southern islands, it is likely that an additional digest of *Tsp45I* can be added after the *NlaIV* digest to distinguish *P. acuta* from *P. damicornis* as indicated in Johnston et al. (2018), although the mtORF haplotypes of *P. damicornis* in French Polynesia are unconfirmed.

## Conclusion

This paper presents and validates a highly accurate, rapid, and low cost sequence-free method for differentiating the cryptic *Pocillopora* species present in Mo’orea and the broader geography of French Polynesia using mtORF and PocHistone amplicons. We demonstrate that RFLP protocols can be developed to identify cryptic *Pocillopora* species in regions other than the Hawaiian Islands; additional protocols could be tailored for other reef regions in the Pacific and the Red Sea (Oury et al., 2025). As cryptic species are increasingly identified across coral genera (Grupstra et al., 2024, Rassmusen et al., 2025), approaches for RFLP protocol development, such as those presented here, are likely to become increasingly necessary globally.

## Supporting information

Cohn et al. Supplemental Materials and Methods

## Author Contributions

The conception of this work is attributed to A.M.S.C., S.C.B., H.M.P., F.M.C., with design by F.M.C. and E.J. under the guidance of the other authors. Data resources (e.g., haplotype alignments) and coral samples were provided by E.J., S.C.B., H.M.P., and P.H. Experiments were performed by F.M.C., K.L., and J.A.S. Data analysis was performed by F.M.C., E.J., S.C.B., and A.M.S.C. The paper primary draft was led by F.M.C. with input from E.J., S.C.B., H.M.P. and A.M.S.C. Figures were generated by F.M.C., J.A.S., E.J., and H.M.P. All authors edited and finalized the manuscript. Funding was acquired by A.M.S.C., S.C.B., and H.M.P.

## Acknowledgements

As guests, we recognize and give thanks for the land and water resources of Polynesia, in particular Mo’orea, and to the traditional owners of the land, both past and present. Māuruuru roa. With respect to the spelling of Tahitian words, we endeavored to follow the Te Fare Vāna’a transcription system that is adhered to by a large segment of the Tahitian community, but also recognize other community members follow the Raapoto transcription system where the island name of Mo’orea is, for example, spelled without the ‘eta (i.e., Moorea). A portion of this research was conducted at the UC Berkeley Gump South Pacific Research Station; we thank the station staff for their support. The samples collected herein are resources of the people and government of French Polynesia. Research was completed under permits issued by the Territorial Government of French Polynesia (Délégation à la Recherche) and the Haut-Commissariat de la République en Polynésie Francaise (DTRT) (MCR LTER Convention d’Accueil 2005–2026, SCB Convention d’Accueil 2019–2023, HMP Convention d’Accueil 2022–2023), and under the Direction de l’environnement (DIREN) permits (HMP N°10626, SCB N°6360). Samples were exported under Convention on International Trade in Endangered Species (CITES) export permits (HMP FR24987000029-E, SCB FR1998700173-E and FR2198700111-E). We thank the Délégation à la Recherche, DTRT, and DIREN for their support. This work represents a contribution of the Mo’orea Coral Reef (MCR) LTER Site.

## Data Accessibility and Benefit-Sharing

### Data Accessibility Statement

All sequence data required for reproduction of these analyses are available on public databases or have been provided as supplemental data, as documented in the main text of the manuscript.

Sanger sequences representing new genetic diversity of the mtORF gene have been deposited in GenBank under accession numbers XXX and XXX. An example protocol with suggested materials and methods for executing this workflow is published (DOI: 10.17504/protocols.io.bp2l6j57rvqe/v1).

### Benefit-Sharing Statement

The results of this research, and all associated data and protocols, are open access and are being shared with the provider communities and the broader scientific community. This research can support conservation and restoration activities related to the organisms being studied - a priority concern in the provider community.

