## Supplementary material for "A Genetic Method for Distinguishing Cryptic *Pocillopora* Species in French Polynesia without Sequencing": Cohn et al. Supplemental Materials and Methods

#### *In Vitro Uniformed Validation*

*Pocillopora* corals were sampled from 2m depth in the lagoon at four separate sites on the north shore of the island of Moorea between 2022-2023 under the French Polynesian Government Délégation à la Recherche (Protocole d'Accueil 2022–2023) and transported under CITES permit FR2398700018-E. Tissue biopsies were stored in DNA/RNA Shield reagent (Zymo Research #R1100-250) at -20°C until DNA extraction. DNAs were extracted using the Quick-DNA 96 Plus Kit (Zymo Research #D4070) following manufacturer's instructions with the addition of proteinase K. Species identification was completed at the University of Rhode Island (URI) following mtORF amplification and Sanger sequencing of the forward primer (FatP6.1; Flot et al. 2008) for *P. acuta*, *P.* cf. *effusa*, *P. tuahiniensis*, and *P. verrucosa*, followed by PocHistone amplification (PocHistoneF and PocHistoneR primers; Flot et al. 2008) and *XhoI* RFLP for *P. meandrina* and *P. grandis* according to Johnston et al. (2018). To confirm the identity of the 36 unknown *Pocillopora* samples used in the uninformed validation (**Supplemental Table S1**), a phylogenetic species tree was generated from their mtORF sequences as well as those of NCBI reference sequences of known haplotype identities **(Supplemental Figure S2).**

#### *BanI/BceAI Digest*

An additional co-digest with *BanI* and *BceAI* is available to differentiate *P.* cf. *effusa* from all other species, and has the added benefit of grouping *P. grandis* (Haplotype 1), *P. meandrina* (Haplotype 1a, 1c, 1d, 1e, 8a, 9), and *P. acuta* (Haplotype 5a) for differentiation with additional digests. *P.* cf. *effusa* shares a fixed adenine SNP at 311 bp with *P. acuta*, *P. meandrina*, and *P. grandis*. However, *P. acuta* and *P. meandrina* share a guanine SNP at 211 bp with all other species except for *P.* cf. *effusa*.

Five microliters of the mtORF amplicon of each species was digested with .5 ul of *BanI* and 0.5 ul of BseAI in 0.5ul of rCutSmart™ Buffer for 1 hour at 37◦C followed by 20 min at 65◦C. The product was then run out on a 2% agarose gel at 100 Volts for 45 min.

All successful digests in **Figure S4** were accurately identified in the uninformed validation using *BanI/BceAI* (substituting for *EcoRV-HF*). However, *BanI/BceAI* was not chosen for the ***Finalized Sequential Protocol*** because it requires two enzymes, instead of the one enzyme required when performing the *EcoRV-HF* digest.

***RADseq Histone Analysis:***

To investigate why some *P. grandis* samples produced a triple banding pattern following digestion of the Pochistone region with the *XhoI* restriction enzyme and determine the phylogenetic placement of the mtORF haplotype 9, we generated an identity by state matrix using restriction site associated DNA sequenced (RADseq) libraries and investigated bam mapping alignments of the PocHistone region using those same libraries (Burgess and Johnston, unpublished data). Samples (<1cm) were collected from 384 *Pocillopora* corals from five species (*P. meandrina*, *P. verrucosa*, *P. grandis*, *P. tuahiniensis*, and *P.* cf. *effusa*) at 5, 10, and 20 m depths from the fore reefs of Moorea in 2019 and 2021. Genomic DNA was extracted from these samples using Omega BIO-TEK E-Z 96 Tissue DNA Kits (D1196-01). Two elutions (50 and 100 uL) were collected in HPLC water and combined. Samples were identified to species prior to restriction site associated DNA sequencing (RADseq) by sequencing the mtORF region and then using the histone RFLP assay to differentiate *P. meandrina* and *P. grandis* following Johnston et al. (2018).

RADseq libraries were prepared by digesting genomic DNA with the DpnII restriction enzyme (New England Biolabs: R0543S). Half reactions of the KAPA HyperPrep Kit (Roche: KK8504) and xGen™ Stubby Adapter and UDI Primers Pairs (Integrated DNA Technologies: 10005921) were used to prepare libraries. DNA fragment size selection was carried out post adapter ligation, pre library amplification, using Mag-Bind® TotalPure NGS beads (Omega BIO-TEK: M1378-01). Equimolar amounts of each library were then pooled and sequenced across four lanes of the HiSeq platform (2 x 150bp).

To identify the placement of mtORF haplotype 9, demultiplexed reads were first mapped to the *P. meandrina* genome (Stephens et al. 2022) using BWA (Li and Durbin 2009) and bam files were filtered with samtools (Li et al. 2009) to remove reads with a mapping quality score below 20 and those with secondary alignments (samtools view -b -q 20 -F 0x0100). Genotype likelihoods for each sample were estimated using ANGSD (v0.940; "-domajorminor 1, -gl 2, -domaf 2, -docounts 1, -dopost 1, -dogeno 16, -doGlf 2, -SNP_pval 1e-7, -minInd 310, -nInd 387, -minmapQ 30, -minQ 30, -doBcf 1, -geno_minDepth 10, -dumpCounts 2, -doIBS 1, -doCov 1, -makeMatrix 1") (Korneliussen et al. 2014) and the resulting identity by state matrix was visualized to identify *Pocillopora* species placement. All haplotype 9 samples grouped with *P. meandrina*.

To investigate why triple banding patterns, instead of the expected double banding pattern, were produced following PocHistone digestion with the XhoI restriction enzyme, RADseq libraries from the nine libraries identified as *P. grandis* in the identity by state matrix were mapped to the PocHistone region (GenBank accession: MG587096: Johnston et al. (2018) using BWA-MEM (Li 2013)). One sample that had been identified as *P. meandrina* by the combination of mtORF sequencing and histone RFLP assay grouped with the *P. grandis* clade in the identity by state matrix. RADseq data from this library was also mapped to the Pochistone region. The Pochistone bam files were then visually inspected in Geneious Prime 2025.1.1. The two libraries that produced the expected double banding pattern for *P. grandis* were found to be homozygous for the restriction enzyme cut site used to identify *P. grandis* (CTCGAG). The six libraries that produced a triple banding pattern for the restriction enzyme cut site used to identify *P. grandis* displayed heterozygosity at the cut site in the bam files (CTCGAG / CTCGCG). The one library that was identified as *P. meandrina* using the RFLP assay, but which grouped with *P. grandis* in the identity by state matrix, was found to be homozygous for the PocHistone site used to identify *P. meandrina* (CTCGCG).

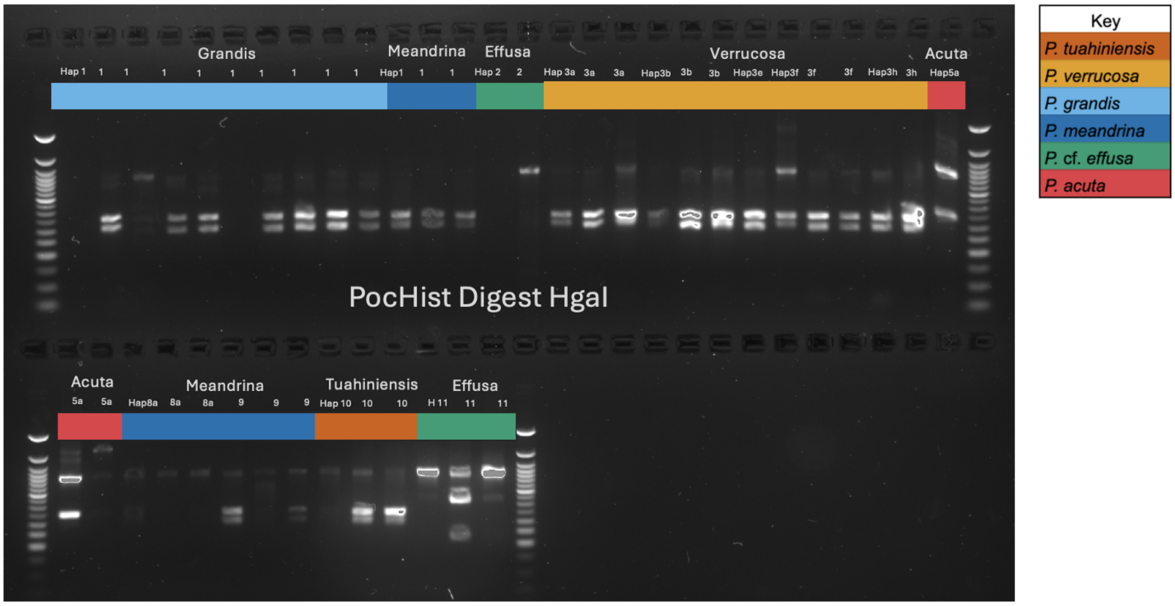


**Figure S1.** Results from an example *in vitro* testing of candidate enzymes on the PocHistone amplicon. The attempted *HgaI* digest was run on the PocHistone amplicon of 42 known *Pocillopora* species representing 12 haplotypes collected in French Polynesia and was designed to differentiate *P.* cf. *effusa* and *P. acuta* from all other species. Digests in lanes 1, 6, and 14 were unsuccessful on this gel. Variability of within species digest pattern (ie. *P.* cf. *effusa*) suggested that this was not a good candidate enzyme to use and further demonstrated the difficulties of using the PocHistone amplicon to differentiate species beyond *P. meandrina* and *P. grandis*.

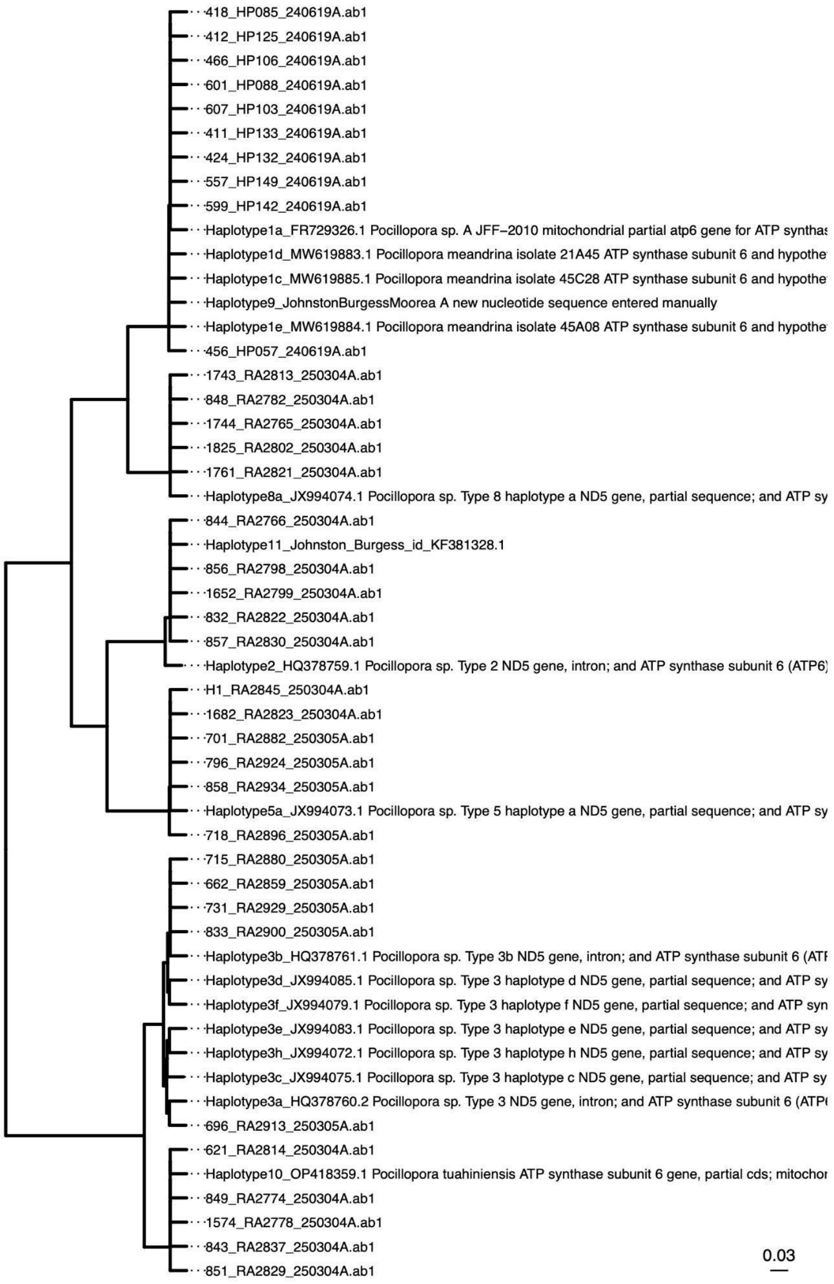


**Figure S2**. Species tree containing GenBank reference sequences of known *Pocillopora* haplotype identities (accession numbers in sequence name), along with sequences of the 36 unknown *Pocillopora* samples (samples ending in ".abi") used in the ***In Silico and In Vitro Uninformed Validation of the Protocol*** **(Supplemental Data S3)**. PocHistone amplicons for these samples were identified prior to the uninformed validation by *XhoI* digest.

### SacI-HF/EcoRV-HF Digest

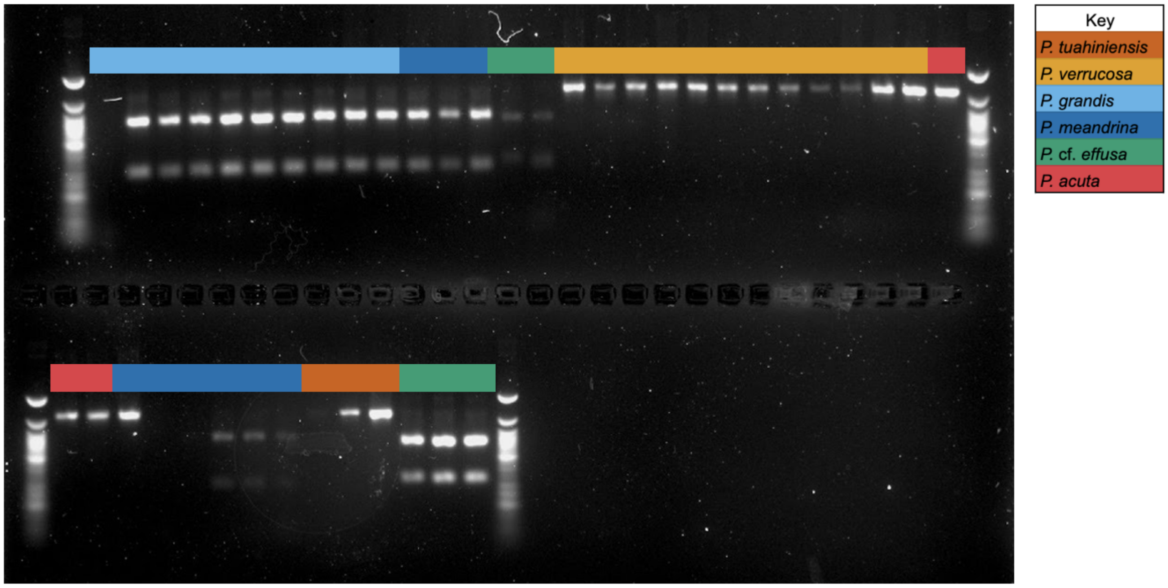


**Figure S3.** Results from initial *in vitro* testing of candidate enzymes. The attempted *SacI-HF/EcoRV-HF* digest was run on the MtORF amplicon of 42 known *Pocillopora* species representing 12 haplotypes (***In Vitro Enzyme Testing Methods***) collected in French Polynesia. Digests in Lanes 1, 32, and 33 were unsuccessful on this gel. This digest was designed to differentiate *P.* cf. *effusa* from all other species while grouping *P. meandrina* with *P. grandis*, and *P verrucosa* with *P. tuahiniensis*. However, *SacI-HF* did not cut haplotype 8a of *P. meandrina*, meaning that haplotype 8a grouped with *P. verrucosa* and *P. acuta* rather than the other *P. meandrina* samples. Therefore it was not selected as a digest for the ***Finalized Sequential Protocol*** because not all haplotypes of the same species produced the same digestion pattern.

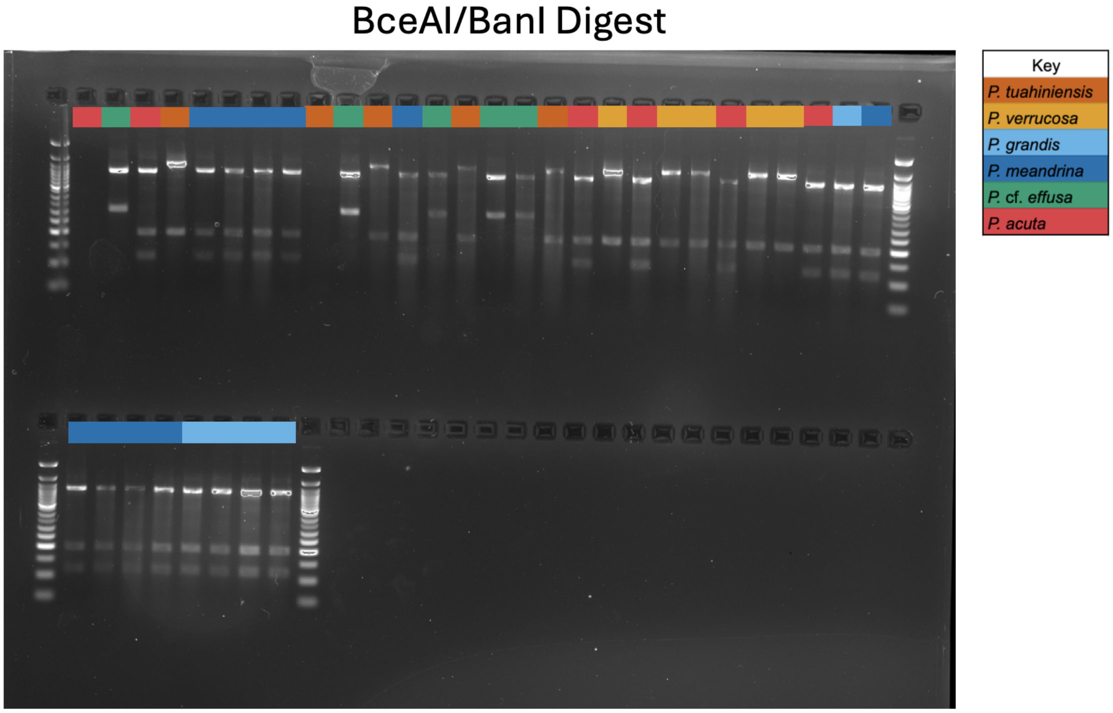


**Figure S4**. Results of the *in vitro uninformed validation* for the *BceAI/BanI* digest run on 36 unidentified Pocillopora samples representing *P. acuta* (n = 6 haplotype 5a), *P. meandrina* (n = 10 haplotypes 1a; 8a), *P.* cf. *effusa* (n = 5 haplotype 11), *P. grandis* (n = 5 haplotype 1a), *P. tuahiniensis* (n = 5 haplotype 10), and *P. verrucosa* ( n=1 haplotype 3a, n=4 haplotype 3b) collected in French Polynesia. Digests in lanes 1 and 9 were unsuccessful on this gel. *BanI* and *BceAI* correctly differentiated *P. cf. effusa* from all other species, and grouped *P. grandis* (Haplotype 1), *P. meandrina* (Haplotype 1a, 1c, 1d, 1e, 8a, 9), and *P. acuta* (Haplotype 5a). However it was not included in the ***Finalized Sequential Protocol*** because *EcoRV-HF* was able to differentiate *P. cf. effusa* with only one enzyme**.**

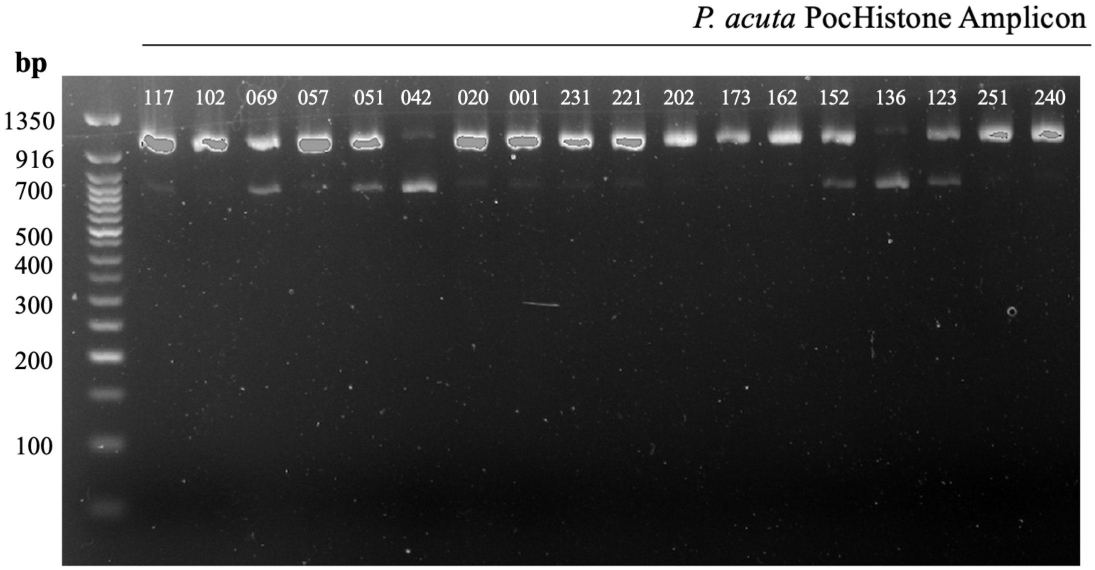


**Figure S5**. Undigested *Pocillopora acuta* PocHistone amplicon showing bands at 1014 bp and 669 bp (or both) depending on whether the sample is likely homozygous or heterozygous for the 345 bp indel in the PocHistone amplicon. Samples from the collection of S. Burgess.

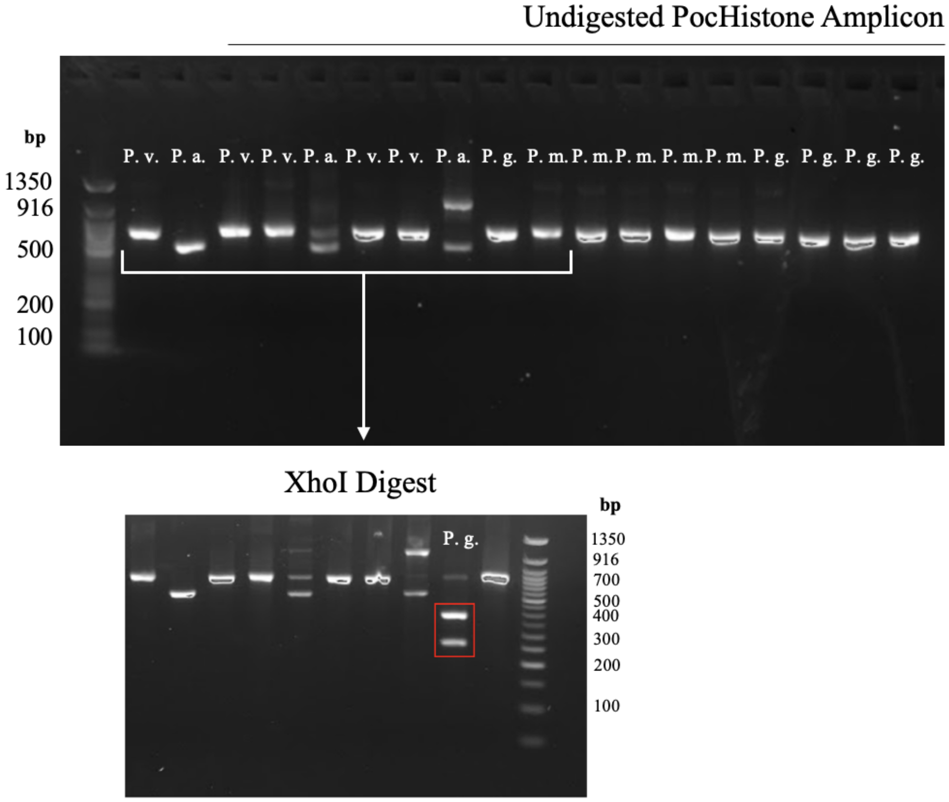


**Figure S6**. Top gel: Undigested PocHistone amplicon for *Pocillopora verrucosa* (P.v.), *Pocillopora acuta* (P.a.), *Pocillopora grandis* (P.g.), and *Pocillopora meandrina* (P.m.). Different band size anomalies are noticeable in the second, fifth, and eight wells - all *P. acuta* samples. Bottom gel: Several of the PocHistone amplicons above are digested with *XhoI*, and despite the anomalies present in the *P. acuta* bands, the *P. grandis* sample (well 9) is the only sample showing the characteristic band sizes of 287 bp and 382 bp that identify it (red box).

| **Date.Collected. dd.mm.yy** | **longitude** | **latitude** | **Sample. ID** | **Species** |
|---|---|---|---|---|
| 9/4/22 | NA | NA | H1 | *P. acuta* |
| 23/03/23 | -149.8526452 | -17.48223933 | 1652 | *P.* cf. *effusa* |
| 23/03/23 | -149.8526452 | -17.48223933 | 1682 | *P. acuta* |
| 23/03/23 | -149.8526452 | -17.48223933 | 1574 | *P. tuahiniensis* |
| 27/03/23 | -149.8526452 | -17.48223933 | 1825 | *P. meandrina* |
| 27/03/23 | -149.8526452 | -17.48223933 | 1744 | *P. meandrina* |
| 27/03/23 | -149.8526452 | -17.48223933 | 1743 | *P. meandrina* |
| 27/03/23 | -149.8526452 | -17.48223933 | 1761 | *P. meandrina* |
| 5/2/23 | -149.8344367 | -17.47810177 | 843 | *P. tuahiniensis* |
| 5/2/23 | -149.8344367 | -17.47810177 | 844 | *P.* cf. *effusa* |
| 5/2/23 | -149.8344367 | -17.47810177 | 849 | *P. tuahiniensis* |
| 5/2/23 | -149.8344367 | -17.47810177 | 848 | *P. meandrina* |
| 5/2/23 | -149.8344367 | -17.47810177 | 856 | *P.* cf. *effusa* |
| 5/2/23 | -149.8344367 | -17.47810177 | 621 | *P. tuahiniensis* |
| 5/2/23 | -149.8344367 | -17.47810177 | 832 | *P.* cf. *effusa* |
| 5/2/23 | -149.8344367 | -17.47810177 | 857 | *P.* cf. *effusa* |

| | | | | |
|---|---|---|---|---|
| 5/2/23 | -149.8344367 | -17.47810177 | 851 | *P. tuahiniensis* |
| 5/2/23 | -149.8344367 | -17.47810177 | 858 | *P. acuta* |
| 4/2/23 | -149.8344367 | -17.47810177 | 715 | *P. verrucosa* |
| 4/2/23 | -149.8344367 | -17.47810177 | 718 | *P. acuta* |
| 4/2/23 | -149.8344367 | -17.47810177 | 696 | *P. verrucosa* |
| 4/2/23 | -149.8344367 | -17.47810177 | 731 | *P. verrucosa* |
| 4/2/23 | -149.8344367 | -17.47810177 | 701 | *P. acuta* |
| 4/2/23 | -149.8344367 | -17.47810177 | 662 | *P. verrucosa* |
| 7/2/23 | -149.8076461 | -17.47569278 | 833 | *P. verrucosa* |
| 7/2/23 | -149.7835691 | -17.47304608 | 796 | *P. acuta* |
| 5/10/22 | -149.8076461 | -17.47569278 | 456 | *P. grandis* |
| 6/10/22 | -149.8076461 | -17.47569278 | 466 | *P. meandrina* |
| 5/10/22 | -149.8076461 | -17.47569278 | 424 | *P. meandrina* |
| 5/10/22 | -149.8076461 | -17.47569278 | 418 | *P. meandrina* |
| 5/10/22 | -149.8076461 | -17.47569278 | 412 | *P. meandrina* |
| 5/10/22 | -149.8076461 | -17.47569278 | 411 | *P. meandrina* |
| 6/10/22 | -149.8076461 | -17.47569278 | 557 | *P. grandis* |
| 5/10/22 | -149.8076461 | -17.47569278 | 599 | *P. grandis* |

| | | | | |
|---|---|---|---|---|
| 6/10/22 | -149.8076461 | -17.47569278 | 607 | *P. grandis* |
| 6/10/22 | -149.8076461 | -17.47569278 | 601 | *P. grandis* |

**Table S1.** Collection and identification information for 36 *Pocillopora* samples (URI) used in the uninformed in vitro identification.

| Species | Haplotypes | Amplicon | RE | Fragment size (bp) | Cuts other Species | Additional Digests | Amplicon | RE | Fragment size (bp) | Cuts other Species |
|---|---|---|---|---|---|---|---|---|---|---|
| *P. verrucosa* | 3a, 3b, 3c, 3d, 3e, 3f, 3h | mtORF | **(1) BseYI<br>C'CCAGC<br>GGGTC'G<br>AciI<br>C'CGC<br>GGC'G** | 209, 338, 430 | All other species<br>430, 546 | **No** | | | | |
| *P. tuahiniensis* | 10 | mtORF | **(1) BseYI<br>C'CCAGC<br>GGGTC'G<br>AciI<br>C'CGC<br>GGC'G** | 145, 186, 209, 430 | All other species<br>430, 546 | **No** | | | | |
| *P. cf. effusa* | 2, 11 | mtORF | **(1) BceAI<br>ACGGC(N)12'<br>TGCCG(N)14'<br>BanI<br>G'GYRCC<br>CCRYG'G** | 311, 663 | mean, grandis, acuta<br>113, 196, 663<br>verr, tuah<br>196, 780 | **No** | | | | |
| *P. acuta* | 5a | mtORF | **(1) NlaIV<br>GGN'NCC<br>CCN'NGG** | 30, 171, 313, 463 | **mean, grandis, cf. effusa<br>202, 313, 463<br>verr, tuah<br>463, 515** | **No** | | | | |
| *P. meandrina* | 1a, 1c, 1d, 1e, 8a, 9 | mtORF | **(1) BceAI<br>ACGGC(N)12'<br>TGCCG(N)14'<br>BanI<br>G'GYRCC<br>CCRYG'G** | 113, 196, 663 | acuta, grandis<br>113, 196, 663<br>verr, tuah<br>196, 780;<br>cf. effusa<br>311, 663 | **(2) NlaIV<br>to remove P. acuta** | (2) PocHistone | **(3) XhoI<br>C'TCGAG<br>GAGCT'C** | 287, 382 (Johnston et al 2018) | All other species<br>669 |

| | | | | | | | | | | |
|---|---|---|---|---|---|---|---|---|---|---|
| *P. grandis* | 1a | mtORF | **(1) BceAI ACGGC(N)12' TGCCG(N)14' BanI G'GYRCC CCRYG'G** | 113, 196, 663 | acuta, grandis 113, 196, 663; verr, tuah 196, 780; cf. effusa 311, 663 | **(2) NlaIV to remove P. acuta** | (2) PocHistone | **(3) XhoI C'TCGAG GAGCT'C** | 669 (Johnston et al 2018) | All other species 669 |

**Table S2**. A summary of an alternative identification scheme replacing the *EcoRV-HF* digest with a *BceAI/BanI*. The table lists each *Pocillopora* species, associated haplotypes, the amplicon and restriction enzyme used for identification, resulting band fragment size of that species, and resulting band fragment size of other species when cut with that restriction enzyme. Additional digests required to fully identify each *Pocillopora* species are also listed.
